# A four-step regulatory cascade controls bistable transfer competence development of the integrative and conjugative element ICE*clc* in *Pseudomonas*

**DOI:** 10.1101/2019.12.20.884213

**Authors:** Nicolas Carraro, Xavier Richard, Sandra Sulser, François Delavat, Christian Mazza, Jan Roelof van der Meer

**Affiliations:** Department of Fundamental Microbiology, University of Lausanne, 1015 Lausanne, Switzerland; Department of Mathematics, University of Fribourg, 1700 Fribourg, Switzerland; UMR CNRS 6286 UFIP, University of Nantes, 44000 Nantes, France

**Keywords:** bistability, integrative and conjugative element, regulation, stochastic modeling, adaptation, feedback control

## Abstract

Genetic bistability controls different phenotypic programs in defined subpopulations of genetically identical bacteria. Conjugative transfer of the integrative and conjugative element ICE*clc* in *Pseudomonas* requires development of a transfer competence state in stationary phase, but this state arises only in 3-5% of individual cells. The mechanisms controlling and underlying the bistable switch between non-active and transfer competence cells have long remained enigmatic. Using a variety of genetic tools combined with stochastic modeling, we characterize here the factors and overall network architecture controlling bistable ICE*clc* activation of transfer competence. Two new key regulators (BisR and BisDC) were uncovered, that link the hierarchical cascade of ICE*clc* transfer competence activation to in total four regulatory nodes. The final activator complex named BisDC drives a positive feedback on its own transcription, and directly controls the “late” ICE promoters for excision and transfer. Stochastic mathematical modeling conceptually explained the arisal and maintenance of bistability by the feedback loop, and demonstrated its importance to guarantee consistent prolonged downstream output in activated cells. A minimized gene set allowing controllable bistable output in a *Pseudomonas putida* in absence of the ICE*clc* largely confirmed model predictions. Phylogenetic analyses further showed that the two new ICE*clc* regulatory factors are widespread among putative ICEs found in *Gamma*- and *Beta*- proteobacteria, highlighting the conceptual importance of our findings for the behaviour of this wide family of conjugative elements.

**Author summary:** Integrative and conjugative elements (ICEs) are mobile genetic elements present in virtually every bacterial species, which can confer adaptive functions to their host, such as antibiotic resistance or xenometabolic pathways. Integrated ICEs maintain by replication along with the genome of their bacterial host, but in order to transfer, the ICE excises and conjugates into a new recipient cell. Single-cell studies on a unique but widely representative ICE model from *Pseudomonas* (ICE*clc*) showed that transfer only occurs from a small dedicated subpopulation of cells that arises during stationary phase conditions. This bistable subpopulation differentiation is highly significant for ICE behaviour and fitness, but how it is regulated has remained largely unknown. The present work unveiled the architecture of the ICE*clc* transfer competence regulation, and showed its widespread occurrence among ICEs of the same family. Stochastic mathematical modeling explained how bistability is generated and maintained, prolonging the capacity of stationary phase cells to complete all stages of ICE activation.

## Introduction

Biological bistability refers to the existence of two mutually exclusive stable states within a population of genetically identical individuals, leading to two distinct phenotypes or developmental programs [1]. The basis for bistability lies in a stochastic regulatory decision resulting in cells following one of two possible specific genetic programs that determine their phenotypic differentiation [2]. Bistability has been considered as a bet-hedging strategy leading to an increased fitness of the genotype by ensuring survival of one of both phenotypes depending on environmental conditions [3]. A number of bistable differentiation programs is well known in microbiology, notably competence formation and sporulation in *Bacillus subtilis* [4,5], colicin production and persistence in *Escherichia coli* [6], virulence of *Acinetobacter baumannii* [7], or the lysogenic/lytic switch of phage lambda [8].

The dual lifestyle of the *Pseudomonas* integrative and conjugative element (ICE) ICE*clc* has also been described as a bistable phenotype (Fig. 1A) [9]. In the majority of cells ICE*clc* is maintained in the integrated state, whereas a small proportion of cells (3-5%) in stationary phase activates the ICE transfer competence program [9–11]. Upon resuming growth, transfer competent (tc) donor cells excise and replicate the ICE [12], which can conjugate to a recipient cell, where the ICE can integrate [10]. ICE*clc* transfer competence comprises a differentiated stable state, because initiated tc cells do not transform back to the ICE-quiescent state. Although tc cells divide a few times, their division is compromised by the ICE and eventually arrests completely [13,14].

**Fig 1.**
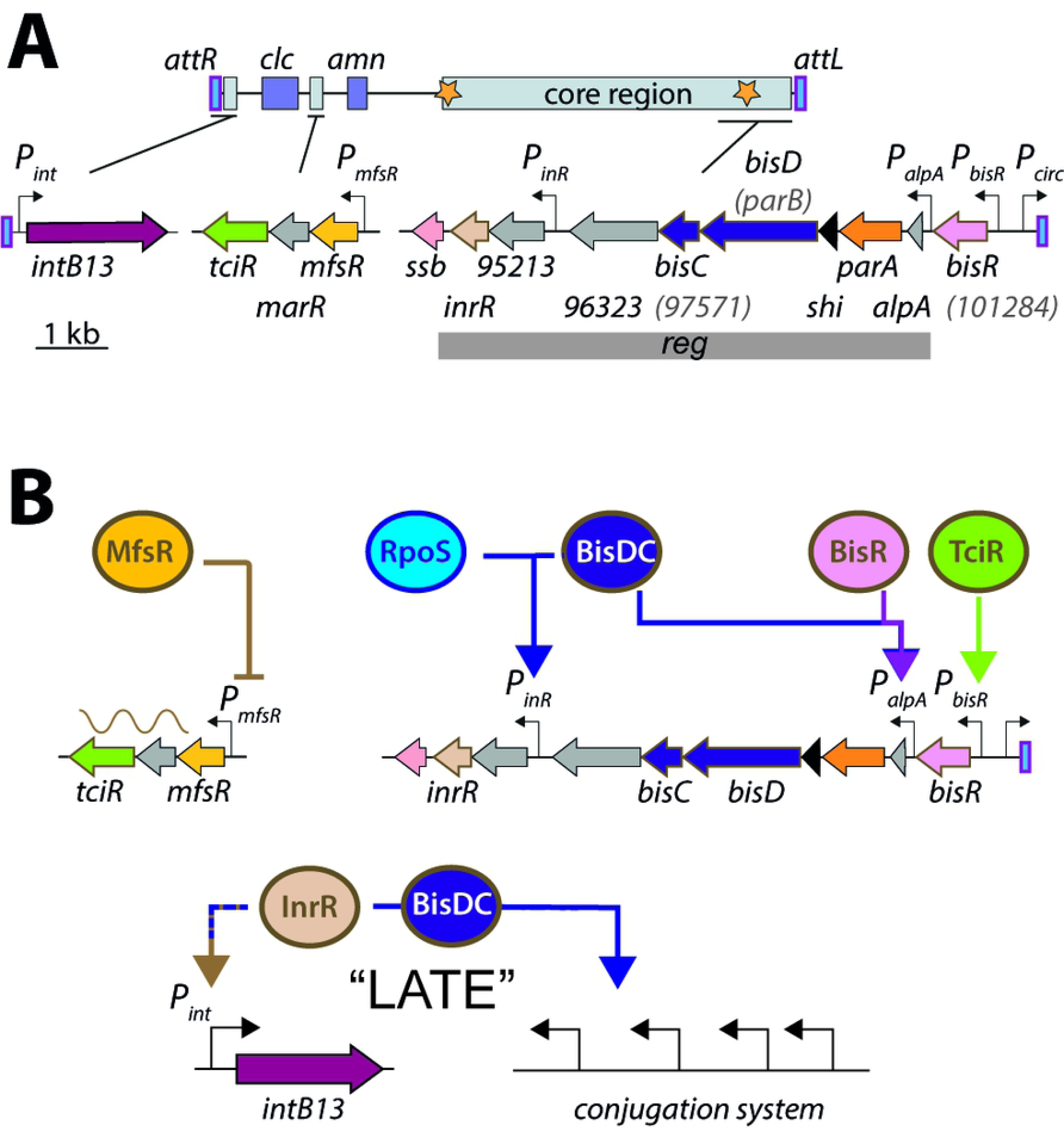
ICE*clc* and its hierarchical regulation network. **A.** Schematic representation of the genetic organization of ICE*clc* (GenBank accession number AJ617740.2). Loci of interest are detailed below the general map and drawn to scale. Genes are represented by colored arrows with their name below (inside parentheses in case of former name). Promoters are represented by hooked arrows pointing towards the transcription orientation. *attL* and *attR*, attachment sites; orange stars, origins of transfer. *clc* genes: 3-chlorocatechol degradation, *amn* genes: 2-aminophenol degradation. **B.** Hierarchy of regulation of ICE*clc* transfer competence (see main text).

ICEs have attracted wide general interest because of the large variety of adaptive functions they can confer to their host, including resistance to multiple antibiotics [15–17], or metabolism of xenobiotic compounds [18,19]. Although the existence and the fitness consequences of the ICE*clc* bistable transfer competence pathway have been studied in quite some detail, the regulatory basis for its activation has remained largely elusive [11]. In terms of its genetic makeup, ICE*clc* is very distinct from the well-known SXT/R391 family of ICEs and from ICE*St1*/ICE*St3* of *Streptococcus thermophilus* [11,20,21]. These carry analogous genetic regulatory circuitry to the lambda prophage, which is characterized by a typical double-negative feedback control [20–22]. Transcriptomic studies indicated that about half of the ICE*clc* coding capacity (~50 kb) is higher expressed in stationary phase cultures grown on 3-chlorobenzoate (3-CBA), and organized in at least half a dozen transcriptional units [23]. A group of three consecutive regulatory genes precludes ICE*clc* activation in exponentially growing cells, with the first gene (*mfsR*) constituting a negative autoregulatory feedback (Fig. 1B) [24]. Overexpression of the most distal of the three genes (*tciR*), leads to a dramatic increase of the proportion of cells activating the ICE*clc* transfer competence program [24]. Despite this initial discovery, however, the nature of the regulatory network architecture leading to bistability and controlling further expression of the ICE*clc* genes in tc cells has remained enigmatic.

The primary goal of this work was to dissect the regulatory control underlying ICE*clc* activation and transfer competence. Our second aim was to understand the conceptual genetic architecture yielding and maintaining ICE bistability. By gene deletions, heterologous expression and mutation studies, we targeted known and unknown regulatory elements encoded by ICE*clc*, and judged their effects on expression of known bistable single-copy ICE*clc* promoter reporter fusions, and on ICE transfer capability. On the basis of a defined subset of key regulators, we developed a mathematical model to stochastically simulate bistability generation in different architectures. We then produced a simplified bistability generator in *P. putida* without ICE*clc* that can yield controllable subpopulation proportions. Our results further indicated that the key ICE*clc* bistability regulatory elements are widespread and conserved, illustrating their importance for the behavior of this conglomerate of related ICEs.

## Results

### A global ICE*clc* activation cascade

Activation of wild-type ICE*clc* in *P. knackmussii* B13 and *P. putida* occurs in stationary phase cells after growth on 3-CBA as sole carbon source [9]. In order to circumvent 3-CBA dependence and permit clearer dissection of the ICE*clc* regulatory network, we chose to express all presumed regulatory factors in *P. putida* strains with or without ICE*clc* from a plasmid-driven IPTG-inducible *P*_*tac*_ promoter, and cultured cells on succinate as sole carbon source (Table 1, Table S1). Transfer of wild-type ICE*clc* from succinate-grown *P. putida* UWC1 to an ICE*clc*-free isogenic *P. putida* tagged with a gentamycin resistance marker was below detection limit, indicating that spontaneous ICE activation under those conditions is negligible (Fig. 2A).

**Table 1.**
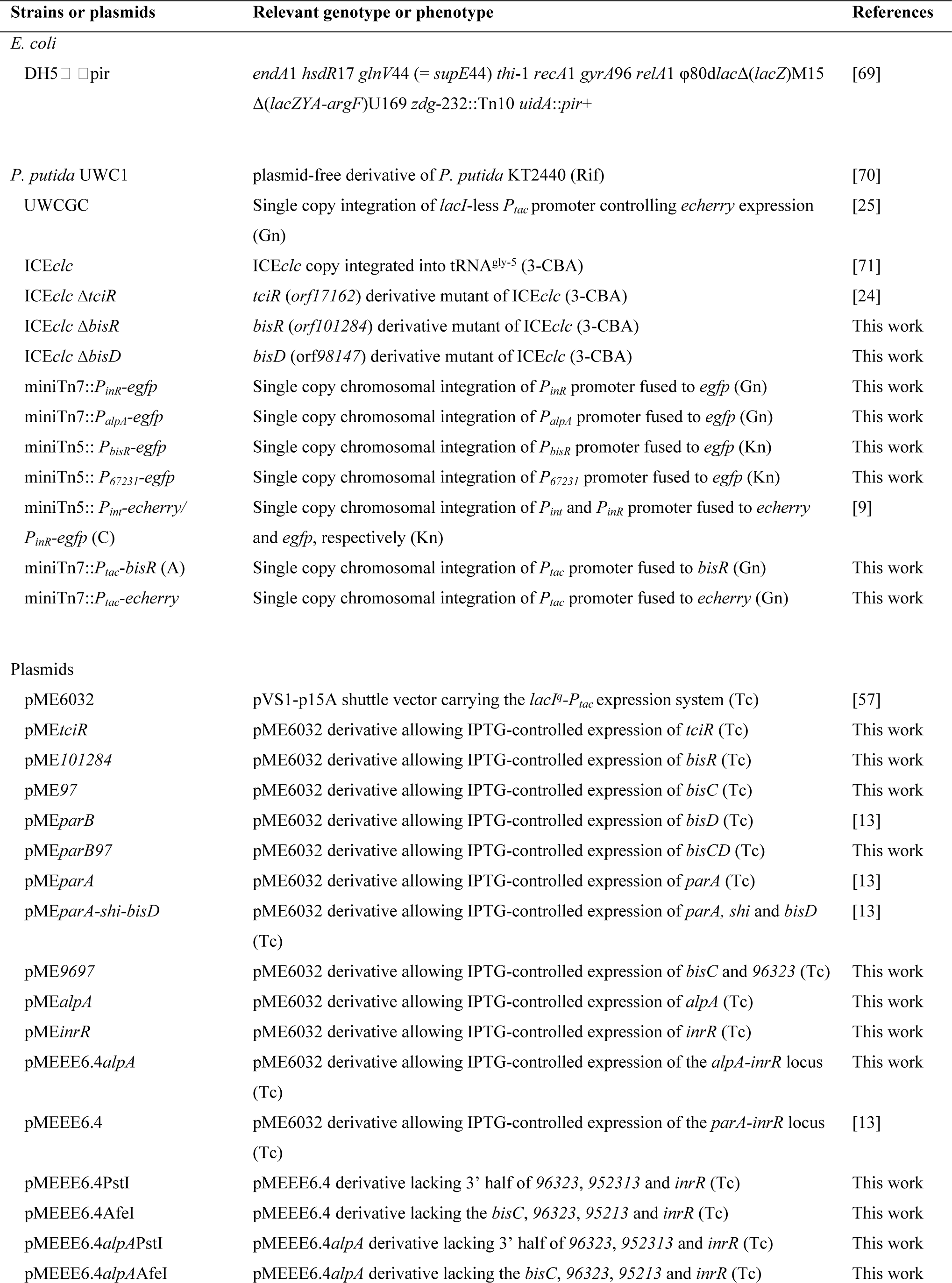
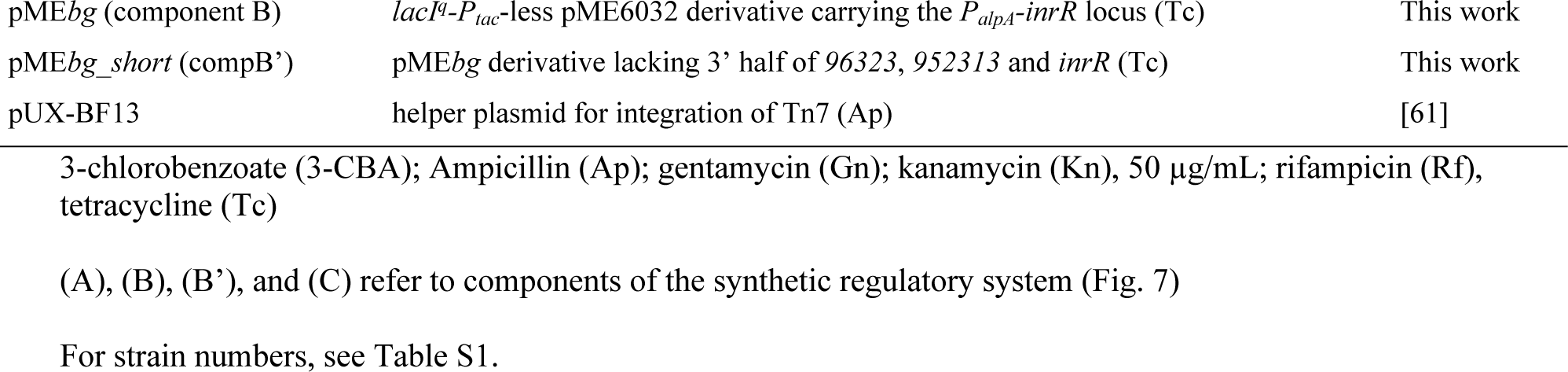
Strains and plasmids used in this study.

**Fig 2.**
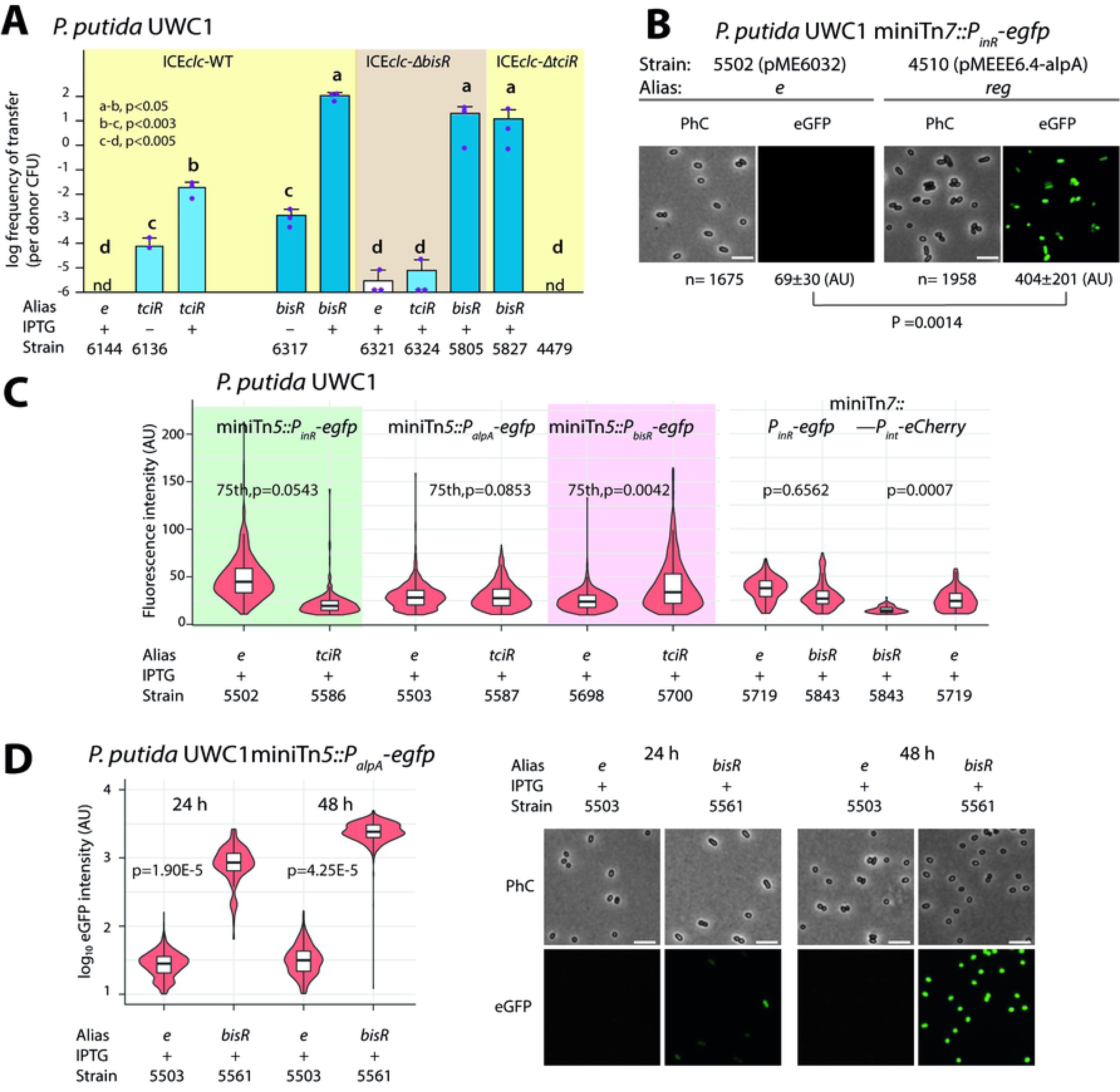
Two regulatory elements link early to late activation of ICE*clc*. **A.** Hierarchical control of *tciR* and *bisR* on ICE*clc* conjugative transfer. Bars show the means of transconjugant formation after 48 h in triplicate matings using *P. putida* UWC1 donors carrying the indicated ICE*clc* or plasmids, in absence (-) or presence (+) of 0.1 mM IPTG, and with *P. putida* UWCGC as recipient. Dots represent individual transfer; nd: not detected (<10^−7^ for the three replicates). Letters indicate significance levels in ANOVA, followed by post-hoc Tukey testing. **B.** IPTG induction (0.05 mM) of a plasmid containing the *reg* fragment of ICE*clc* in *P. putida* (pMEEE6.4*alpA*) is sufficient to drive expression of *egfp* from a single-copy inserted *P*_*inR*_-promoter, which is otherwise silent (*e*, pME6032). **C.** Reporter expression from single copy *P*_*bisR*_, *P*_*inR*_, *P*_*int*_, or *P*_*alpA*_ fusions in *P. putida* UWC1, in presence of the empty plasmid pME6032 (e), pMEtciR (*tciR*), or pME*bisR* (*bisR*). Diagrams show representative fluorescence distributions (violin plots) in 24- or 48-h cultures grown on succinate, induced with 0.05 mM IPTG. Note how TciR only activates *P*_*bisR*_. p-values calculated in t-tests on triplicate values of the 75^th^ percentile, given the skewed distributions. **D.** Activation of the *P*_*alpA*_ promoter by BisR. Diagram shows the ^10^log reporter fluorescence in *P. putida* expressing *bisR* (pME*bisR*) or not (*e*, pME6032), grown for 24 h on succinate medium, induced with 0.05 mM IPTG. P-values calculated in t-tests on triplicates of the median fluorescence. Selected microscopy images show the strong eGFP fluorescence in individual *P. putida* UWC1 cells from the single-copy miniTn5::*P_alpA_-egfp* reporter in case of *bisR* induction. PhC, phase-contrast. eGFP, eGFP fluorescence (artificially colored to green). Images scaled to same absolute minimum and maximum intensity (for **B** and **D**).

Previous work had unveiled an operon on ICE*clc* of three consecutive regulatory genes consisting in a transcriptional repressor named *mfsR*, *marR* and a gene coding for a LysR-type activator named *tciR* (Fig. 1A) [24]. Increasing induced expression of *tciR* from *P*_*tac*_ on a medium copy plasmid (pME*tciR*) in *P. putida* UWC1-ICE*clc* was sufficient to trigger ICE*clc* transfer from succinate-grown cells up to frequencies close to the one observed after growth on 3-CBA [25] (10^−2^ transconjugant colony-forming units (CFU) per donor CFU, Fig. 2A; strain 6136). On the other hand, induction of *tciR* was insufficient to trigger eGFP production from the known ICE*clc* bistable promoter *P*_*inR*_ in absence of the ICE (Fig. 2C, strain 5586). These results confirmed that TciR is a central activator for ICE*clc* transfer but implied the existence of a further regulatory cascade linking TciR to the known ICE*clc* bistable promoters [24,26].

Earlier work had suggested the implication of two genes located on a 6.4-kb fragment at the leftmost end of ICE*clc* (close to the *attL* site) in regulating the *P*_*int*_-integrase promoter (Fig. 1A) [27]. One of these was the gene coding for the protein InrR [7], the other was a gene with an annotated ParB domain (Fig. 1A) [27]. This fragment was cloned separately and elongated to encompass the *alpA* gene (Fig. 1A, *reg* fragment), since a promoter had been mapped upstream of *alpA* [23]. IPTG induction of *P. putida* UWC1 without ICE*clc* carrying a plasmid with this 6.6-kb *reg* fragment under control of *P*_*tac*_ (pMEEE6.4*alpA*) resulted in activation of eGFP from a single copy chromosomally-integrated *P*_*inR*_-*egfp* reporter, which otherwise is completely silent (Fig. 2B). This indicated that one or more regulatory functions were encoded on the 6.6-kb *reg* fragment.

Next, we tested the hypothesis that TciR directly or indirectly activates the native promoter upstream of *alpA*. Induction of *tciR* from *P*_*tac*_ on pME*tciR* in *P. putida* UWC1 containing a single-copy *P*_*alpA*_-*egfp* transcriptional fusion did not yield any eGFP fluorescence (Fig. 2C, strain 5587). As we had previously noticed that there is another open reading frame upstream of *alpA* named *orf101284*, which forms an independent transcriptional unit (Fig. 1A) [23], we tested whether induction of *tciR* could activate the promoter of this gene. Indeed, IPTG-induction of *tciR* in *P. putida* UWC1 with a single-copy *P*_*101284*_-*egfp* reporter (renamed to *P*_*bisR*_-*egfp*) yielded weak but significant eGFP fluorescence compared to the same *P. putida* host with the empty vector (Fig. 2C, strain 5700; p=0.0042 on means of 75th percentiles from biological triplicates). This suggested that the link between TciR and further ICE activation proceeds through transcription of *orf101284.* We renamed this open reading frame to *bisR*, or <u>bis</u>tability regulator, for its presumed implication in ICE*clc* bistability control (Fig. 1B, see further below).

### BisR is a new regulator crucial for relaying ICE*clc* transfer competence activation

*bisR* is predicted to encode a 251-aa protein of unknown function with no detectable Pfam-domain. Further structural analysis using Phyre2 suggested three putative domains with low confidence (between 38% and 53%) (Fig. S1) [28]. One of these is a predicted DNA-binding domain, which hinted at the possible function of BisR as a transcriptional regulator. BlastP analysis showed that BisR homologs are widely distributed and well conserved among *Beta-, Alpha*- and *Gammaproteobacteria*, with homologies ranging from 43–100% amino acid identity over the (quasi) full sequence length (Fig. S2).

In order to investigate its potential regulatory function, *bisR* was cloned under the control of the *P*_*tac*_ promoter on pME*bisR* and introduced into *P. putida* UWC1-ICE*clc*. Increasing induction of *bisR* from *P*_*tac*_ triggered high rates of ICE*clc* transfer on succinate media (Fig. 2A, strain 6317). Deletion of *bisR* on ICE*clc* abolished its transfer (strain 6321), even upon overexpression of *tciR* (strain 6324), but could be restored upon ectopic expression of *bisR* (Fig. 2A, strain 5805). This showed that the absence of transfer was due to the lack of intact *bisR*, and not to a polar effect of *bisR* deletion on the downstream gene *alpA* (Fig. 1A). In addition, transfer of an ICE*clc* deleted for *tciR* [24] could be restored by ectopic *bisR* expression (Fig. 2A, strain 5827). This indicated that TciR is ‘upstream’ in the regulatory cascade of BisR, and that TciR does not act anywhere else on the expression of components crucial for ICE*clc* transfer.

Despite triggering ICE*clc* transfer, single-copy reporter transcriptional fusions showed no direct activation of *P*_*int*_ or *P*_*inR*_ by BisR (Fig. 2C). This suggested that BisR was an(other) intermediate regulator step in the complete cascade of activation of ICE*clc* transfer competence. Among different tested known ICE–promoters, BisR only triggered very strong expression from a single copy *P*_*alpA*_–*egfp* transcriptional fusion (Fig. 2D). This linked BisR as the intermediate regulator between TciR and the 6.6-kb *reg* fragment (Fig. 1B).

### A new regulator BisDC is the last step in the activation cascade

In order to identify the master activator(s) controlling expression of the late genes of ICE*clc*, the *reg* fragment was subcloned in different gene and promoter configurations (Fig. 3A). Similar as for the *P*_*inR*_–promoter (Fig. 2B), IPTG induction of the full *reg* fragment (strain 5721) triggered expression of *echerry* from *P*_*int*_ in *P. putida* without ICE*clc* (Fig. 3B). Removing *alpA* had no measurable effect on *P*_*int*_–*echerry* expression (Fig. 3B, pMEEE6.4, strain 5720), but *echerry* expression was absent when the *P*_*tac*_ promoter was replaced by the natural promoter *P*_*alpA*_ (Fig. 3B, pME*bg*, strain 5985). Removing an internal region covering the 3’ extremity of *orf96323*, *orf95213* as well as the 5’ extremity of *inrR* (Fig. 3A, pMEEE6.4*alpA*PstI, strain 6045), still yielded expression of the *P*_*int*_–*echerry* reporter, albeit less strong (Fig. 3B). In contrast, a plasmid encompassing only *parA-shi-bisD* and the 5’ extremity of *bisC* (pMEEE6.4*alpA∆*AfeI, Fig. 3B, strain 6044), was unable to activate the *P*_*int*_–reporter. This pointed to the importance of having the *bisD* and/or *bisC* genes for *P*_*int*_– expression. Neither induction of *bisC*, of *bisD* or of *inrR* alone led to reporter expression (Fig. 3B, strains 5724, 5884 or 5776). However, when *bisC* was coordinately induced from *P*_*tac*_ with the upstream gene *bisD*, *echerry* was expressed from *P*_*int*_ in *P. putida* UWC1 without ICE*clc* (Fig. 3B, pME*parB97*, strain 6059). These results indicated that BisDC acts as an ensemble to activate the bistable *P*_*int*_–promoter and pointed to *bisDC* as the last step in the transcription activator cascade of ICE*clc*. Pfam analysis detected a DUF2857-domain in the BisC protein, and further structural analysis using Phyre2 indicated significant similarities of BisC to FlhC (Fig. S1) [28]. FlhC is a subunit of the master flagellar activator FlhCD of *E. coli* and *Salmonella* [29,30]. BisD (formerly named ParB-like protein) carries a ParB domain, with a predicted DNA binding domain in the C-terminal portion of the protein (Fig. S1). Although, no FlhD domain was detected in BisD, in analogy to FhlCD we named the ICE*clc* activator complex BisDC, for <u>bis</u>tability regulator subunits <u>D</u> and <u>C</u>.

**Fig 3.**
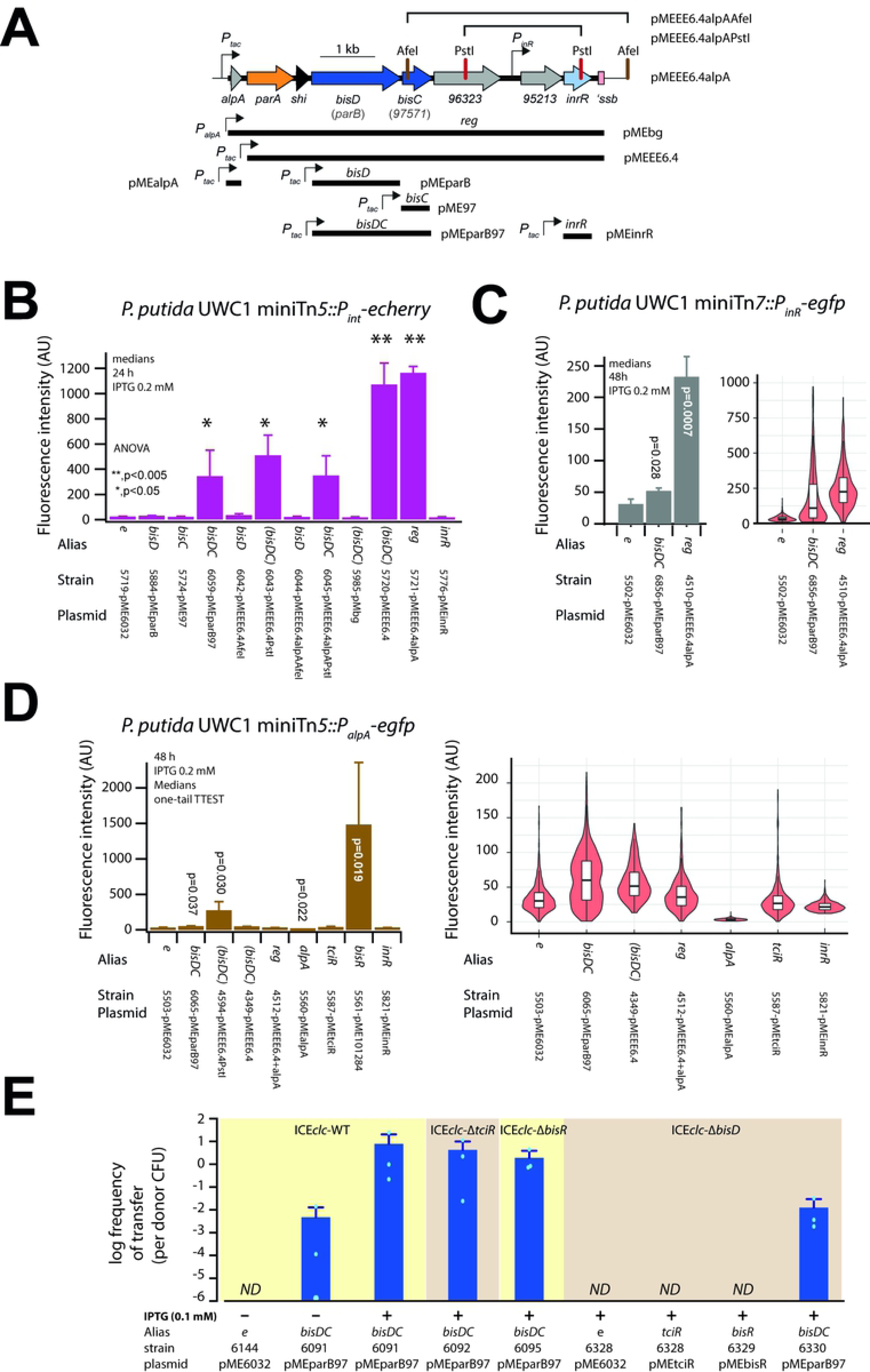
BisCD is the master activator complex of ICE*clc*. **A.** Schematic representation of ICE*clc* main regulatory locus (drawn to scale) and indications of the cloned regions in pME6032 to test promoter activation. Plasmid names as in the other panels and in main text. **B.** Activation of eCherry fluorescence from single-copy integrated *P*_*int*_ promoter fusion in *P. putida* without ICE*clc*, complemented with the various regulatory plasmid constructs of A. Bars represent the mean of median fluorescence in biological triplicates (±SD), measured on individual cells after 24 h growth in succinate medium and induced with IPTG (0.05 mM). Stars indicate significance levels in ANOVA followed by post-hoc Tukey test. **C.** As **B**, but on the *P*_*inR*_-promoter with *P. putida* cells imaged after 48 h. p-values obtained from paired t-test comparison of triplicate medians between isogenic *P. putida* carrying empty pME6032 and the indicated plasmid constructs. Note how induction of *bisDC* from plasmid pME*parB97* yields a highly skewed eGFP expression in comparison to pMEEE6.4*alpA* (violin plots). **D.** As **B**, but showing activation of *P*_*alpA*_ by BisDC. **E.** Effect of *bisDC* induction on conjugative transfer of ICE*clc* wild-type or mutant derivatives. Transfer assay details as in legend to Figure 2.

IPTG induction of *bisDC* from plasmid pME*parB97* was also sufficient to activate the previously characterized bistable *P*_*inR*_-promoter in *P. putida* UWC1 (Fig. 3C, p=0.028). The distribution of eGFP-values among induced cells was highly skewed (Fig. 3C, violon plots, *bisDC*), whereas expression of eGFP from *P*_*inR*_ was more homogenous in *P. putida* UWC1 expressing the *reg* fragment (pMEEE6.4*alpA*) (Fig. 3C). Expression of eGFP from *P*_*inR*_ in a double-labeled single-locus-integrated reporter strain was weaker than that of *echerry* from *P*_*int*_ and higher when plasmid fragments did not include *inrR* itself (Fig. S3, e.g., pMEEE6.4PstI vs pMEEE6.4), although induction of *inrR* alone was insufficient to yield *P*_*int*_-*echerry* expression (Fig. 3B). To better understand this, we revisited activation of the *P*_*alpA*_-*egfp* reporter in *P. putida* without ICE*clc*. Induction of *bisDC* alone also yielded significant eGFP fluorescence from *P*_*alpA*_ (Fig. 3D, strain 6065), but not as high as by induction of the fragment encompassed by plasmid pMEEE6.4PstI (Fig. 3A, strain 4594), and in comparison to induction by BisR (Fig. 3D, strain 5561). Induction of *inrR* alone had no influence on *P*_*alpA*_-*egfp* reporter expression (Fig. 3D, strain 5821). In contrast, induction of *alpA* alone significantly lowered expression from *P*_*alpA*_ compared to an empty plasmid control (Fig. 3D, strain 5560). Co-induction of *alpA* also lowered the magnitude of BisDC activation of *P*_*alpA*_ (Fig. 3D, pMEEE6.4, strain 4349 vs pMEEE6.4*alpA*, strain 4512). These results indicated that BisDC can activate *P*_*alpA*_, whereas AlpA may have a modulary repressive role. Since *P*_*alpA*_ is the only mapped promoter upstream of the *reg* fragment (Fig. 3A), this would imply that BisDC controls its own expression.

Increasing induction of *bisDC* from plasmid pME*parB97* again yielded high frequencies of ICE*clc* transfer from *P. putida* UWC1 under succinate-growth conditions (Fig. 3E, strain 6091). Expression of BisCD also induced transfer of ICE*clc* deleted for *tciR* or for *bisR* (Fig. 3E). This confirmed that both *tciR* and *bisR* relay activation steps to *P*_*bisR*_ and *P*_*alpA*_, respectively, but not to further downstream ICE promoters (Fig. 1B). Moreover, an ICE*clc* deleted for *bisD* could not be restored for transfer by overexpression of *tciR* (strain 6328) or *bisR* (strain 6329), but only by complementation with *bisDC* induction (Fig. 3E, strain 6330). Interestingly, the frequency of transfer of an ICE*clc* lacking *bisD* complemented by expression of *bisDC* in *trans* was two orders of magnitude lower than that of similarly complemented wild-type ICE*clc*, of ICE*clc* deleted for *tciR* or for *bisR* (Fig. 3E). This result suggested the necessity of a ‘reinforcement’ occurring in the wild-type configuration that was lacking in the *bisD* deletion and could not be restored by *in trans bisDC* induction.

### BisDC creates a positive feedback loop that maintains ICE*clc* bistable output

Having identified the main regulatory elements in the ICE*clc* activation cascade, we next aimed to determine the origin of bistable activation [9]. First, we tested the influence of overexpressing each of the identified regulatory elements individually by IPTG induction in *P. putida* UWC1 containing ICE*clc* on the activation of a single copy insertion of a dual *P*_*int*_– and *P*_*inR*_–fluorescent reporter under succinate-growth conditions. IPTG-induction of *tciR* from plasmid pME*tciR* yielded clear bimodal output of individual cells expressing eGFP and mCherry (Fig. 4, Table S2). In contrast, IPTG-induction of *bisR* from pME*bisR* led to eGFP and mCherry fluorescence in all cells in a unimodal fashion (Fig. 4). Induction of either *bisD* or *bisC* alone resulted again in bimodal output but smaller subpopulation sizes (Fig. 4). In contrast, induction of *bisD* and *bisC* simultaneously yielded high unimodal fluorescence of both reporters in all cells (Fig. 4). These results suggested that the arisal of bistability is inherent to the architecture of the activation cascade and is dependent on the expression levels of the individual regulatory nodes.

**Fig 4.**
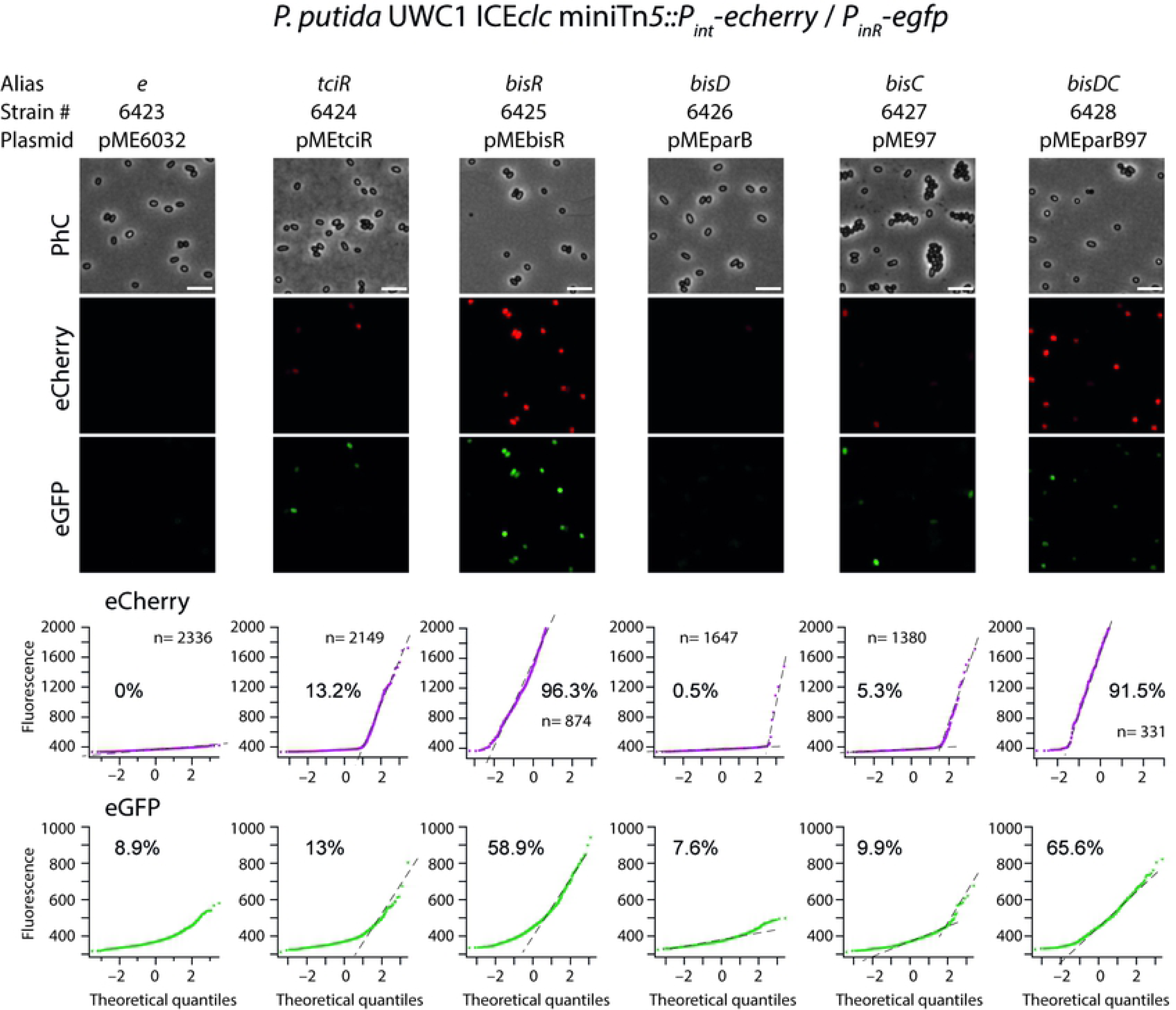
Effect of overexpression of ICE*clc* regulators on subpopulation size of transfer competent cells. Panels show representative images in phase-contrast (PhC) or fluorescence channels (scaled to same minima and maxima) of reporter expression in *P. putida* UWC1-ICE*clc* with an integrated single copy double fusion of *echerry* to *P*_*int*_ and *egfp* to *P*_*inR*_, and complemented with the indicated plasmid. Cells imaged after 48 h growth in succinate minimal medium and induced with IPTG (0.05 mM). Diagrams below images show quantile-quantile plots of measured versus expected normal fluorescence intensity distribution among *n* measured individual cells. Dotted lines indicate the inferred separation of the second (sub)population with the estimated percentage of cells in that subpopulation. Note how QQ plotting can only be applied for subpopulation estimations when these are proportionally small [12,65]. Consequently, overexpressed *bisR* and *bisDC* lead to expression in all cells. White bar on PhC images represents 5 µm.

To better understand how the architecture of the ICE*clc* regulatory factors would generate bistability, we developed a conceptual stochastic mathematical model (Fig. 5A, SI model). The model takes into account production and degradation rates of TciR, BisR and BisDC, their oligomerization, as well as their binding to and unbinding from their respective linked promoters (*P*_*bisR*_, *P*_*alpA*_, Fig. 1B). Of note that model parameters were non-empirical and conceptually imposed. Given their likely modulatory roles, we did not include AlpA or InrR in the model. Stochastic simulations (n=10,000 traces) of the barest feedback loop (BisDC activating *P*_*alpA*_, no TciR or BisR, Fig. 5A), yielded a population with two BisDC output states after 100 time steps, one of which is *zero* and the other with a mean *positive* BisDC value (Fig. 5B). The output *zero* results when BisDC levels stochastically fall to 0 (as in Fig. 5A), since in that case there is no BisDC to stimulate its own production (note that the simulation is arbitrarily started with a binomial distribution with a mean of 8 BisDC, Fig. 5B). Parameter variation indicated that the proportion of output *zero* from the loop is dependent on the binding and unbinding constants for the *alpA* promoter, and the BisDC degradation rate (Fig. 5A, B, different *A1*, *A2* and *A4-*values).

**Fig 5.**
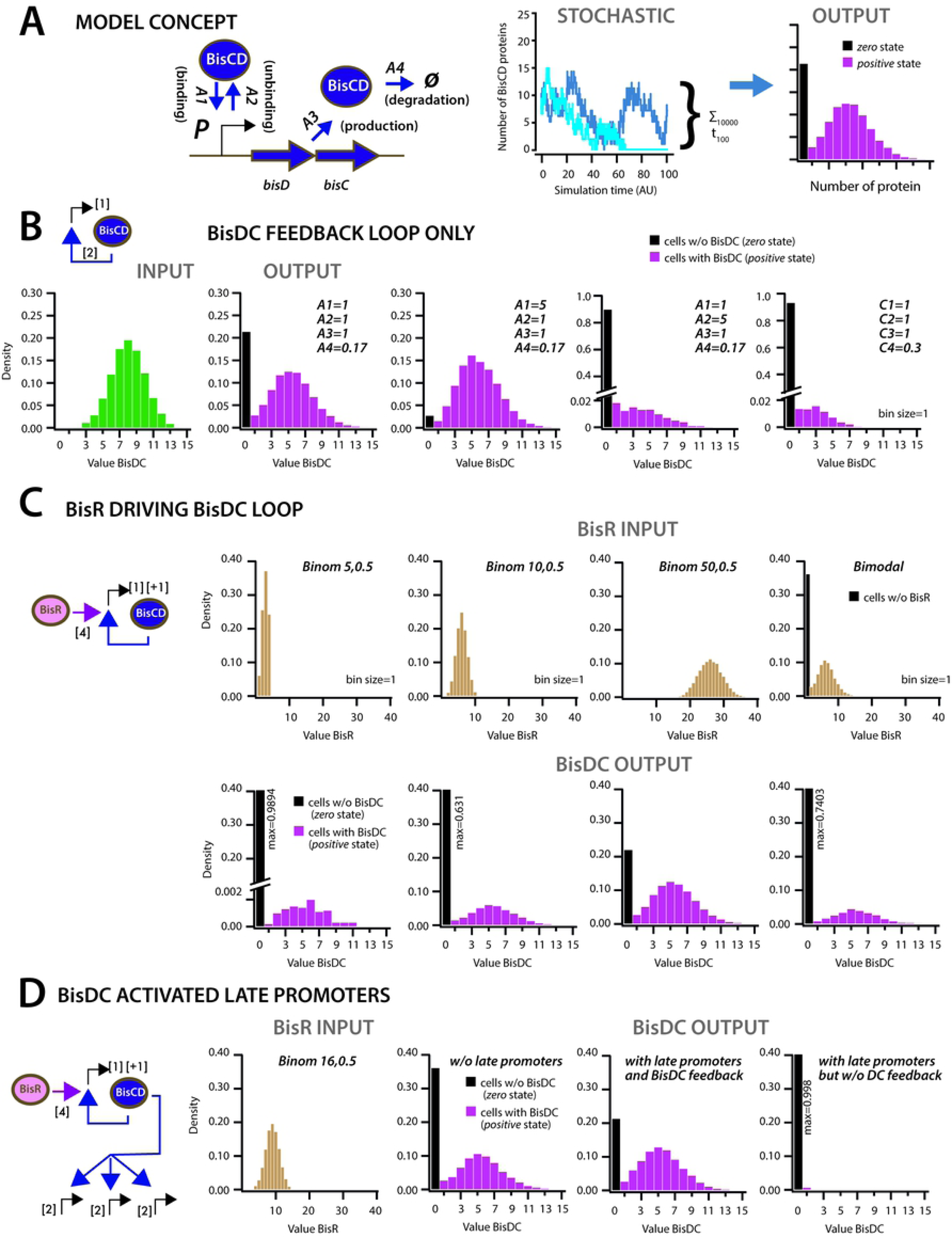
BisDC-sufficient autoregulatory feedback to yield bistable output. **A.** Conceptual stochastic modeling of BisDC protein levels after 100 time steps among 10,000 stochastic simulations of the indicated gene network architecture and parameter configurations. **B.** Behaviour of a BisDC autoregulatory feedback loop on BisDC levels as a function of different binding (*A1*), unbinding (*A2*), production (*A3*) and degradation (*A4*) rate constants, starting from a uniformly distributed set of BisDC levels (input, in green). Model assumes two BisDC proteins needed to activate production of itself (simplified network cartoon corresponds to more elaborate model in A, numbers within brackets indicate interacting or produced protein numbers). Black bar indicates the proportion of cells with *zero* output (i.e., non-activated circuit). **C.** As for A, but in case of BisR-tetramer initiating *bisDC* expression, with different input distributions (uniformly low to high, or bimodal BisR input). Note how higher or bimodal BisR input is not expected to change the BisDC distribution in active cells, but only the proportion of ‘cells’ with *positive* (magenta, bars) and *zero* state (black bars). **D.** As for B, but assuming a further three target downstream promoters titrating each two BisDC proteins, and in presence or absence of the BisDC feedback loop. Note how in absence of BisDC feedback the proportion of cells in *positive* state drastically decreases.

In presence of BisR, the production of BisDC can be initiated from *P*_*alpA*_ by BisR as well as become reinforced by produced BisDC itself (Fig. 5C). Upon a single pulse of BisR, one can see how the feedback loop again ensures BisDC maintenance and leads to a bistable population with *zero* and a mean positive BisDC levels (Fig. 5C). Increasing the input level of BisR results in lowering the proportion of cells with *zero* BisDC state (Fig. 5C). In contrast to the BisDC loop alone, therefore, activation by BisR only influences the proportion of *zero* and *positive* BisDC states, whereas the mean *positive* BisDC output remains the same (Fig. 5C). Interestingly, even a bimodal BisR input leads to a bimodal BisCD output, but to a higher proportion of *zero* BisDC state (Fig. 5C, bimodal). Upon including three other BisDC target promoters (e.g., late promoters such as *P*_*int*_, Fig. 1B), the mean amount of BisDC in *positive* cells remains the same in presence of the BisDC feedback loop. In absence of a feedback loop, however, there are basically no cells with *positive* BisDC after 100 time steps (Fig. 5D).

Appearance of BisR can be connected in the model by the input of TciR (Fig. 6A), in which case a stochastic timely development and subsequent maintenance of BisR and BisDC appear that in some instances lead to a *zero* BisR and thus *zero* BisDC state (Fig. 6B). Depending on the relative amounts of TciR in cells, the proportions of *zero* and *positive* states of both BisR and BisDC vary (Fig. 6C), reminescent of the ICE*clc* wild-type and the ∆*mfsR* mutant (in which case *tciR* is overexpressed, leading to massively higher proportions of tc cells [24]). The effects of bistable state are propagated to downstream functions, as illustrated in Fig. 6D. In presence of the feedback loop, protein output from BisDC regulated promoters is higher and more consistently present in the *positive* cell population, and much lower in its absence (Fig. 6D). This suggests that the first important function of the loop is to ensure sufficient BisDC levels to activate downstream promoters within the *positive* cell population, and, secondly, to stabilize stochastic (input) expression of TciR and BisR to a consistent subpopulation of cells with *positive* BisDC state (Fig. 6C). Cells with *positive* BisDC state represent those that develop ICE*clc* transfer competence.

**Fig 6.**
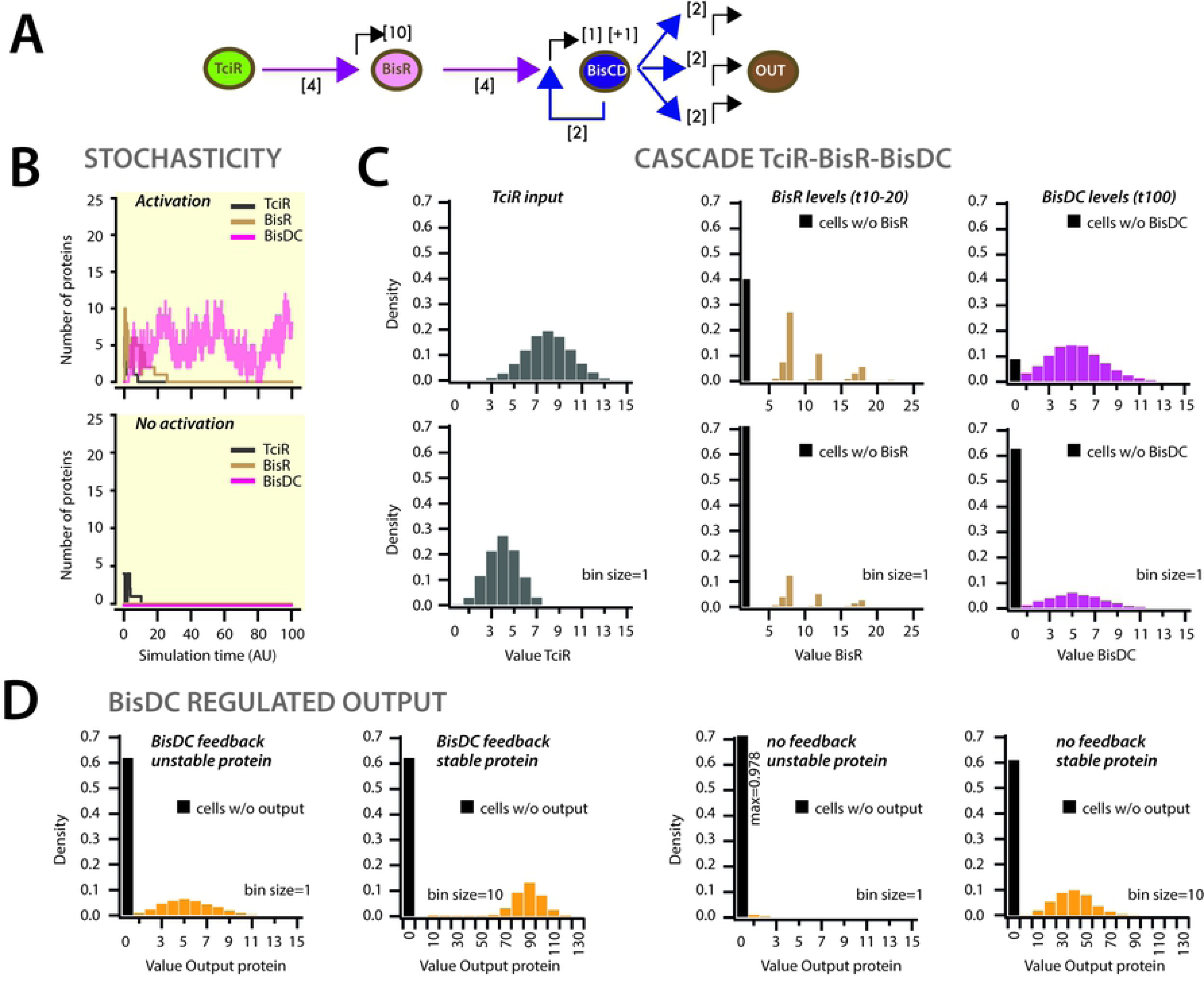
Simulation of the ICE*clc* hierarchical regulatory cascade. **A.** Conceptual representation of the ICE*clc* transfer competence regulation cascade, in which TciR-tetramer (hence the value of 4 in the cartoon link) is assumed to activate *bisR* expression (output=10), with a BisR-tetramer activating *bisDC* expression (output=1), and BisDC-dimer controlling its own and late promoter expression. **B.** Depending on the TciR starting levels, stochastic fluctuations (single trajectories shown across 100 time steps) lead to BisR and BisDC activation and maintenance, or to *zero* state (lower panel). **C.** Regulatory factor level distributions from two different TciR starting distributions across 10,000 simulations; for BisR integrated between time points 10 and 20) and for BisDC after 100 time steps. Bimodal expression of *zero* and *positive* states arises at the *bisR* node, but is further maintained to constant BisDC output as a result of the feedback loop. **D.** In absence of BisDC-feedback, output of a BisDC-dependent protein expression strongly decreases, dependent on the assumed protein degradation constant.

### A minimal inducible bistable system from ICE*clc* components

To further validate the model predictions *in vivo*, we engineered a minimal bistable system with three components. The first component (A) consisted in *bisR* under IPTG-inducible control of *P*_*tac*_, which allows production of BisR in a unimodal manner regardless the concentration of IPTG (Fig. 7A, Fig. S4). The second component consisted in the feedback loop, produced either by a long gene fragment starting from *P*_*alpA*_ and encompassing all genes from *alpA* to *inrR* (B) or a shorter fragment from *alpA* until *bisDC* (B’). The third component (C) contained the dual *P*_*int*_-*echerry* and *P*_*inR*_-*egfp* reporter to measure the BisDC system output (Fig. 7A). Combinations of the different components introduced into *P. putida* without ICE*clc* confirmed that neither B or B’ alone, nor A alone could activate the reporters (Fig. 7B). In contrast, combinations of either A and B, or A and B’ led to activation of the reporters, but only in presence of IPTG to kickstart BisR production (Fig. 3B and 7B). Increasing induction of BisR led to increased proportions of cells with induced reporters (Fig. 7C), as expected from model predictions (Fig. 7D), but not their average fluorescence (Fig. 7E).

**Fig 7.**
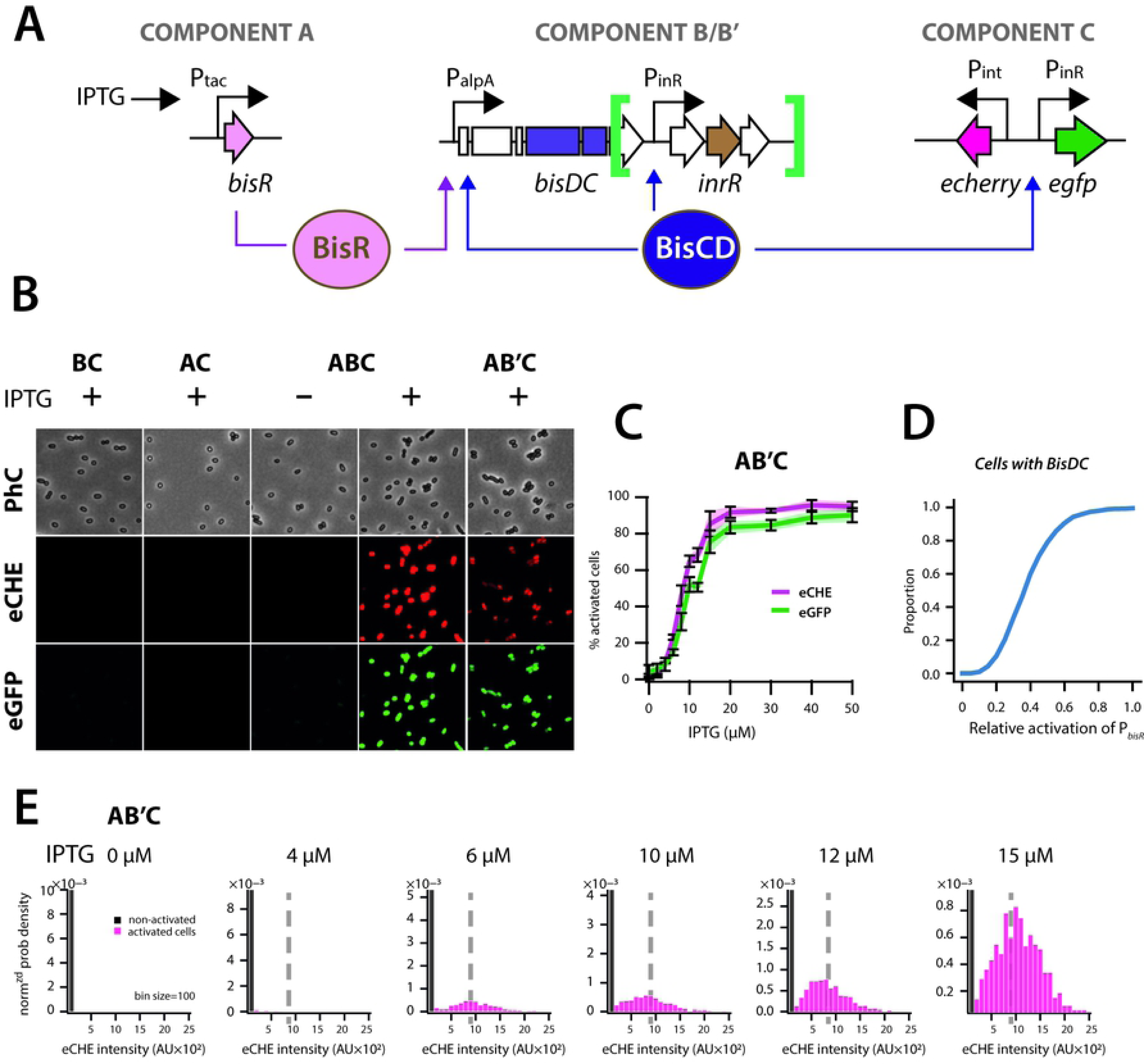
Reconstruction of a minimized inducible ICE*clc* bistability generator. **A.** Schematic representation of the three components used to reconstruct ICE*clc*-based bistability in *P. putida* UWC1 devoid of the ICE. **B.** Imaged output of individual *P. putida* cells carrying the different components as indicated, after growth for 48 h (component B) or 24 h (component B’) in succinate minimal medium, in presence or absence of IPTG (0.1 mM). Images scaled to same minima and maxima, but artificially colored red and green. **C.** Percentage of active cells from QQ-plotting for both reporters in the AB’C configuration after 24 h in succinate minimal medium as a function of IPTG concentration. Lines correspond to the means from three biological replicates with transparent areas and error bars representing the standard deviation. **D.** Calculated proportion of cells expressing *positive* state of BisDC in the model of Figure 5C as a function of their mean BisR starting levels. **E.** Measured distributions of eCherry fluorescence values among the subpopulations of activated (determined from QQ-plotting) and non-active cells at different IPTG concentrations, showing same subpopulation median output (dotted lines of eCherry intensity), as predicted from the feedback model.

### BisCD-elements are widespread in other presumed ICE

Database searches showed that *bisDC* loci are widespread among pathogenic and environmental *Gamma*- and *Beta-proteobacteria*, and are also found in some *Alphaproteobacteria* (Fig. S2). Phylogenetic analysis using the more distantly related sequence from *Dickeya zeae* MS2 as an outgroup indicated several clear clades, encompassing notably *bisDC* homologs within genomes of *Pseudomonas aeruginosa* and *Xanthomonas* (Fig. S5). Several genomes contained more than one *bisDC* homolog, the most extreme case being *Bordetella petrii* DSM12804 with up to four homologs belonging to four different clades (Fig. S5).

Gene synteny from *bisR-bisDC* to *inrR*-*ssb* was maintained in several genomes (Fig. S2), suggesting them being part of related integrated ICEs. Notable exceptions included a region in *P. aeruginosa* Carb01_63, which carried an integrase gene upstream of *bisR* but that was still downstream of a *tRNA*^*gly*^ gene (Fig. S2). This region may encompass an ICE that has retained the same integration specificity as ICE*clc* but carrying a different modular architecture where regulation and integration modules are next to each other, instead of at the opposite ends as in ICE*clc* (Fig. 1 and S2). Further exceptions included genomic regions in *P. aeruginosa* strains HS9 and W60856, which carry a gene for a LysR-type transcriptional regulator (LTTR) upstream of *bisR* with 53% and 51% amino acid identity to TciR (overlap lengths 95%), respectively [31]. Regulation of these two elements might thus involve a *cis*-acting LTTR, rather than the *trans-*acting TciR. In the genomes of *Xanthomonas campestris* strain AW13 and *Cupriavidus nantongensis* X1 (Fig. S2), *bisD* homologs are actually split in two individual genes, one coding for a canonical-length ParB and the other for a BisD-homolog with only DNA-binding domain, suggesting that *bisD* on ICE*clc* may have arisen from a gene fusion.

## Discussion

Understanding regulation of bistability is of fundamental interest for bacterial differentiation and adaptation processes. In this work, we characterized the different regulatory elements controlling bistable activation of the transfer competence program of ICE*clc*. Our results revealed that the previously observed bistable ICE*clc* activation in a minority of cells in a population in stationary phase [9], originates in a positive feedback loop that is maintained by a novel but widespread regulatory complex named BisDC (Fig. 1B, 5, 6). Genetic dissection and expression of individual components in *P. putida* devoid of the ICE showed that the feedback loop is initiated from the successive action of regulatory factors, composed of MfsR, TciR and BisR (Fig. 1B). The previously characterized *mfsR-marR-tciR* operon [24], whose transcription is controlled through autorepression of MfsR, is probably the main break on activation of ICE*clc* in exponential phase, as deletion of *mfsR* results in overexpression of TciR and leads to massively increased ICE transfer [24]. We showed here that TciR activates solely the transcription of a hitherto unrecognized transcription factor gene named *bisR*, and no further critical ICE*clc* promoters (Fig. 2). The BisR amino acid sequence revealed only very weak homology to known functional domains, thus making it the prototype of a new family of transcriptional regulators (Fig. S1). BisR on its turn only activated a single other target, the *alpA* promoter, stimulating production of (among others) BisCD (Fig. 2). The genes *bisD* and *bisC* code for subunits of an activator complex that weakly resembles the known regulator of flagellar synthesis FlhCD, and related regulator complexes controlling activation of other mobile genetic elements [20,32–34]. Finally, we showed that BisDC is sufficient to activate the previously characterized bistable ICE*clc* promoters *P*_*int*_ and *P*_*inR*_. Importantly, we found that BisDC also activates transcription from *P*_*alpA*_, thereby creating a positive feedback loop on its own expression. Our conceptual mathematical model suggested that this feedback loop can both create and maintain the bistable output, although bimodality may already arise at the level of BisR expression and be continued by the feedback architecture (Fig. 5, 6).

Bistable gene network architectures are characterized by the fact that expression variation is not resulting in a single mean phenotype, but can lead to two (or more) stable phenotypes -mostly resulting in individual cells displaying either one or the other phenotype [35–37]. Importantly, such bistable states are an epigenetic result of the network functioning and do not involve modifications or mutations on the DNA [38,39]. Bistable phenotypes may endure for a particular time in individual cells and their offspring, or erode over time as a result of cell division or other mechanism, after which the ground state of the network reappears. One can thus distinguish different steps in a bistable network: (i) the bistability *switch* that is at the origin of producing the different states, (ii) a *propagation* or *maintenance* mechanism and (iii) a *degradation* mechanism [11].

Some of the most well characterized bistable processes in bacteria include competence formation and sporulation in *Bacillus subtilis* [37]. Differentiation of vegetative cells into spores only takes place when nutrients become scarce or environmental conditions deteriorate [40,41]. Sporulation is controlled by a set of feedback loops and protein phosphorylations, which culminate in levels of the key regulator SpoOA~P being high enough to activate the sporulation genes [37]. In contrast, bistable competence formation in *B. subtilis* is generated by feedback transcription control from the major competence regulator ComK. Stochastic variations among ComK levels in individual cells, ComK degradation and inhibition by ComS, and noise at the *comK* promoter determine the onset of *comK* transcription, which then reinforces itself because of the feedback mechanism [42,43]. Maintenance of the ICE*clc* transfer competence pathway thus resembles DNA transformation competence in *B. subtilis* in its architecture of an auto-feedback loop (BisDC vs ComK). However, the switches leading to bistability are different, with ICE*clc* depending on a hierarchy of transcription factors (MfsR, TciR and BisR), and transformation competence being a balance of ComK degradation and inhibition of such degradation [42,43]. ICE*clc* bistability architecture is clearly different from the well-known double negative feedback control exerted by, e.g., the phage lambda lysogeny/lytic phase decision in *E. coli* [8,44,45]. That switch entails essentially a balance of the counteracting transcription factors CI, CII and Cro [46,47]. Interestingly, other ICEs of the SXT/R391 family carry this typical double negative feedback loop architecture, which may therefore control their (bistable) activation [20,22,32].

Mathematically speaking, the ICE*clc* transfer competence regulatory architecture has two states, one of which is *zero* (inactive) and the other with a *positive* value (activation of transfer competence, Fig. 5, 6). Stochastic modeling suggested that the feedback loop maintains *positive* output during a longer time period than in its absence (although it will drop to *zero* at infinite time). Previous experimental data suggested that the tc cells indeed do not return to a silent ICE*clc* state, but become irreparably damaged, arrest their division and wither [13]. However, because their number is proportionally low, there is no fitness cost on the population carrying the ICE [48,49]. The advantage of prolonged feedback loop output is that constant levels of the BisCD regulator can be maintained, allowing coordinated and organized production of the components necessary for the ICE*clc* transfer itself. This would consist of, for example, the relaxosome complex responsible for DNA processing at the origin(s) of transfer, and the mating pore formation complex [10,25]. Because *Pseudomonas* cells activate ICE*clc* transfer competence upon entry in stationary phase, the feedback loop may have a critical role to ensure faithful completion of the transfer competence pathway during this period of limiting nutrients, and to allow the ICE to react as soon as nutrients become again available to cells. Interestingly, ICE*clc* excises in donor cells only when new nutrients become available and cell exit stationary phase [50].

Although our results were relatively conclusive on the roles of the key regulatory factors (MfsR, TciR, BisR, BisDC), there may be further auxiliary and modulary factors, and environmental cues that influence the transfer competence network. For example, we previously found that deletions in the gene *inrR* drastically decreased ICE*clc* transfer capability by 45–fold and reduced reporter gene expression from *P*_*int*_ [9]. Expression of InrR alone, however, did not show any direct activation of *P*_*int*_, *P*_*inR*_ or *P*_*alpA*_ (Fig. 3), and InrR is thus unlikely to be a direct transcription activator protein. On the other hand, effects of inducing cloned ICE*clc* fragments carrying or not *inrR* did affect expression of those three promoters in a *P. putida* without ICE*clc* (Fig. 3). Our results also indicated that induction of AlpA may repress output from the *P*_*alpA*_ promoter, suggesting a further possible modulation of the feedback loop that is initiated by BisR and maintained by BisDC. Previous results also highlighted the implication of RpoS for *P*_*inR*_ activation (Fig. 1B), which may be more generally important for other ICE*clc* regulatory promoters as well [50].

Our results suggested that it might be primarily the levels of transcription factors at different time points in cells, which determine the onset and extent of transfer competence activation (Fig. 4). Exponentially growing cells of both *P. putida* and *P. knackmussii* B13 containing ICE*clc* show no signs of transfer competent cells, which we have attributed to MfsR autorepression of *tciR* transcription [24]. Since we could so far not find any ligand specifically derepressing MfsR [24,51], we assume that basal production of TciR in exponentially growing cells is simply too low or its degradation too fast to activate the next node (the *bisR* promoter, as simulated in the model of Fig. 6B). Cells in stationary phase and particularly those that grew on 3-CBA would then contain sufficient TciR to cause activation of *P*_*bisR*_. We acknowledge, however, that we have so far not measured TciR concentrations and changes thereof in exponentially or stationary phase *P. putida* ICE*clc* cells. Neither do we know whether TciR in wild-type situation associates with a chemical ligand [52] for the subsequent activation of *bisR* transcription. BisR, in contrast, was a very proficient activator of the *alpA* promoter, and in a concentration dependent manner (Figs. 2, 4, 7), suggesting that it is solely the BisR levels in the cells that determine activation of the BisCD bistable feedback loop.

Phylogenetic analyses showed the different ICE*clc* regulatory loci (i.e., *bisR-alpA-bisDC-inrR*, Fig. 1A) to be widely conserved in Beta- and Gammaproteobacteria (Fig. S2, S4). Most likely, these regions are part of ICE*clc*-like elements in these organisms, several of which have been detected previously [18]. The results from our study suggest that they may all have the same core control of their transfer competence regulation. The ICE-like elements in *Bordetella petrii* DSM12804 and at least one ICE*clc*-like element in *P. aeruginosa* JB2 were shown to excise from the chromosome, indicating that they are functional [26,53]. Although the *bisR-alpA-bisDC-inrR* locus was very conserved both in sequence and gene synteny, some other configurations were found (Fig. S2), such as genes coding for LTTR regulators immediately upstream of *bisR* in *P. aeruginosa* strains HS9 (2) and W60856. These may play a role in ICE activation control different from the one exerted by TciR in ICE*clc*.

In conclusion, we dissected the major regulatory hierarchy of the ICE*clc* bistable transfer competence network. *Inter alia*, we also showed how ICE bistability can be regenerated from a minimized system, which might be interesting for applications where bistable phenotypes need to be engineered in clonal bacterial populations [54].

## Material and methods

### Strains and growth conditions

Bacterial strains and plasmid constructions used in this study are shortly described in Table 1 and with more detail in Table S1. Strains were routinely grown in lysogeny broth (LB Miller, Lab Logistics Group) at 30°C for *P. putida* and 37°C for *E. coli* in an orbital shaker incubator, and were preserved at −80°C in LB broth containing 15% (vol/vol) glycerol. Reporter assays and transfer experiments were performed in minimal media [55] supplemented with 10 mM sodium succinate or 5 mM of 3-chlorobenzoate (3-CBA). Antibiotics were used at the following concentrations: ampicillin (Ap), 100 µg/mL for *E. coli* and 500 µg/mL for *P. putida*; gentamycin (Gn), 10 µg/mL for *E. coli*, 20 µg/mL for *P. putida*; kanamycin (Kn), 50 µg/mL; tetracycline (Tc), 12 µg/mL for *E. coli*, 100 µg/mL or 12.5 µg/ml for *P. putida* grown in LB and minimal media, respectively. For induction, bacterial cultures were supplemented with isopropyl β-D-1-thiogalactopyranoside (IPTG) at indicated concentration.

### Molecular biology methods

Plasmid DNA was purified using the Nucleospin Plasmid kit (Macherey-Nagel) according to manufacturer’s instructions. All enzymes used in this study were purchased from New England Biolabs. PCR reactions were carried out with primers described in Table S2. PCR products were purified using Nucleospin Gel and PCR Clean-up kits (Macherey-Nagel) according to manufacturer’s instructions. *E. coli* and *P. putida* were transformed by electroporation as described by Dower *et al*. [56] in a Bio-Rad GenePulser Xcell apparatus set at 25 µF, 200 V and 2.5 kV for *E. coli* and 2.2 kV for *P. putida* using 2-mm gap electroporation cuvettes (Cellprojects). All constructs were verified by DNA sequencing (Eurofins).

### Plasmids and strains construction

Different ICE*clc* gene configurations were cloned in *P. putida* using the broad host-range vector pME6032, allowing IPTG-controlled expression from the LacI^q^-*P*_*tac*_ promoter [57]. Genes *tciR, bisR, bisC (97571), bisCD (parB-97571), bisC+96323* (*97571+96323*), *alpA* and *inrR* were amplified using primer pairs tciREcorI.F/tciREcorI.R, 101284EcorI.F/101284EcorI.R, 97571EcoRI.F/97571RBSEcorI.R, parBEcorI.F/97571EcoRI.F, 96323EcorI.F/97571RBSEcorI.R, alpAEcoRI.F/alpAEcoRI.R, and inrREcorI.F/inrREcorI.R respectively (Table S3), and using genomic DNA of *P. putida* UWC1-ICE*clc* as template. Amplicons were digested by EcoRI and cloned into EcoRI-digested pME6032 using T4 DNA ligase, generating pME*tciR*, pME*bisR* and pME*97*, pME*parB97*, pME*9697*, pME*alpA*, and pME*inrR*. pMEEE6.4 carries the 6.4 kb fragment from pTCB177 containing ICE*clc* locus from *parA* to *inrR* [13,27]. The *alpA*-gene was added to pMEEE6.4 by first PCR amplification of an *alpA-parA-shi-‘parB* fragment using primer pair KpnI_98147_For/ EcoRI_100952_Rev2, KpnI and EcorI digestion of the resulting fragment and cloning into pME6032 digested with the same enzymes. The resulting plasmid was digested with SalI, and the 4.8-kb fragment containing the *P*_*tac*_ promoter, *alpA-parA-shi* and the beginning of *parB* was recovered and used to replace the *parA-shi-parB* part of pMEEE6.4 in order to generate pMEEE6.4*alpA*. pMEEE6.4PstI/pMEEE6.4*alpA*PstI *and* pMEEE6.4AfeI/pMEEE6.4*alpA*AfeI were respectively generated by PstI and AfeI digestion of pMEEE6.4 or pMEEE6.4*alpA*, purification of the high molecular weight product by gel extraction kit using PCR Clean-up kit (Macherey-Nagel), and subsequent ligation using T4 DNA ligase.

The promoter region upstream of *bisR* was amplified using primer pairs Fw_101284_(BamHI)/Rev_101284_(XbaI), digested by BamHI and XbaI, and cloned into pBAM1 [58] digested by the same enzymes to produce pBAM1(miniTn*5*::*P*_*bisR*_-*egfp*). The *alpA* promoter was amplified using primer pair 101’169fw/101’428rv, digested by BamHI and PstI, and subsequently cloned in front of *egfp* into pUC18-derived miniTn*7* delivery plasmid digested by the same enzyme. The *P_inR_-egfp* insert was recovered from the miniTn*5*-based reporter system [9] using HindIII and KpnI, and subsequently cloned into pUC18miniTn*7* digested by the same enzymes. The resulting inserts were integrated in single copy into the chromosomal site a*ttB*_Tn7_ of *P. putida* by using pUX-BF13 for miniTn*7*, or randomly for miniTn*5*-based constructs [58–61], in which case three independent clones were recovered, stored and analyzed.

Component B/B’ plasmids for regeneration of ICE*clc* bistability derive from pMEEE6.4, which was first digested using PmlI and BamHI in order to remove the portion containing *P*_*tac*_, *lacI*, *parA*, *shi* and part of *bisD*. A new insert containing *P_alpA_, alpA, parA, shi* and the missing part of *bisD* was synthesized (ThermoFisher Scientific), and ligated using a Quick-Fusion cloning kit (Bimake). Component B’ was then generated by PstI digestion to remove the part downstream of *bisD* and religated. Deletion mutants in ICE*clc* were constructed using the two-step chromosomal gene inactivation technique as described elsewhere [50,62] and primers listed in Table S2.

### ICE*clc* transfer assays

ICE*clc* transfer was tested with 24h-succinate-grown donor and recipient cultures. Cells were harvested by centrifugation of 1 ml (donor) or 2 ml (recipient, gentamycin-resistant *P. putida* UWCGC) for 3 min at 1200×*g*, washed in 1 ml of minimal media (MM) solution without carbon substrate, centrifuged again and finally resuspended in 20 µl of MM. Donor or recipient alone, and a donor-recipient mixture were deposited on 0.2–µm cellulose acetate filters (Sartorius) placed on MM agar plates, and incubated at 30°C for 48 hours. The cells were recovered from the plates in 1 ml of MM and serially diluted before plating. Donors, recipients and exconjugants were selected on minimal media agar plates containing appropriate antibiotics and/or carbon source (3-CBA).

### Molecular phylogenetic analysis

BisDC phylogeny was inferred from 148 aligned homolog sequences by using the Maximum Likelihood method based on the Tamura-Nei model [63], eliminating positions with less than 95% site coverage. The final dataset was aligned using MEGA7 [64] and contained a total of 2091 positions. Initial tree(s) for the heuristic search were obtained automatically by applying Neighbor-Joining and BioNJ algorithms to a matrix of pairwise distances estimated using the Maximum Composite Likelihood (MCL) approach, and then selecting the topology with superior log likelihood value.

### Fluorescent reporter assays

For the detection of eGFP and eCherry in single cells, *P. putida* strains were cultured overnight at 30 °C in LB medium. The overnight culture was diluted 200 fold in 8 mL of MM supplemented with succinate (10 mM) and appropriate antibiotic(s), and grown at 30 °C and 180 rpm for 24, 48, and 72 hours after inoculation. At indicated sampling times, 150 µl of culture were taken, vortexed for 30 seconds at max speed, and drops of 5 µl were deposited on a regular microscope glass slide (VWR) coated with 1% agarose-in-MM. Cells were covered with a 24 ×50 mm cover slip (Menzel-Gläser) and imaged immediately with a Zeiss Axioplan II microscope equipped with a 100× Plan Achromat oil objective lens (Carl Zeiss), and a SOLA SE light engine (Lumencor). A SPOT Xplorer slow-can charge coupled device camera (1.4 Megapixels monochrome w/o IR; Diagnostic Instruments) fixed on the microscope was used to capture images. Up to ten images at different positions were acquired using Visiview software (Visitron systems GMbH), with exposures set to 40 ms, 500 ms and 500 ms for phase contrast (PhC), eGFP and eCherry, respectively. Cells were automatically segmented on image sets using pipelines described previously [12,48], from which their fluorescence (either eGFP or eCherry, or both) was quantified. Subpopulations of transfer competent cells were quantified using quantile-quantile-plotting as described previously [12,65].

### Statistical analysis

Fluorescent reporter intensities were compared among biological triplicates. In case of mini-Tn*5* insertions, this involved three clones with potentially different insertion sites, each measured individually. For mini-Tn*7* inserted reporter constructs, we measured three biological replicates of a unique clone. Expression differences between mutants and a strain with the same genetic background but carrying the empty pME6032 plasmid were tested on triplicate means of individual median values in a one-sided t-test (hypothesis being that the mutant expression is higher than the control), or in case of extremely skewed population distributions, on the mean of 75^th^ percentiles. Coherent simultaneous data series were tested for significance of reporter expression differences in ANOVA, followed by a post-hoc Tukey test. Violin plots were produced using *ggplot2* in R or Graphpad Prism 8 (www.graphpad.com).

### Mathematical model of ICE*clc* activation

ICE*clc* activation was simulated as a series of stochastic events in different biochemical reaction network configurations (as schematically depicted in Figs. 5 and 6, SI model). TciR, BisR, BisDC and protein output levels were then simulated using the Gillespie algorithm [66,67], implemented in *Julia* using its DifferentialEquations.jl package [68]. 10,000 individual simulations were conducted per network configuration during 100 time steps, during or after which the remaining protein levels were counted and summarized. The code for the mathematical implementation is provided in the Supplementary Informations.

## Acknowledgments

The authors thank Fabrize Merz and Noëmie Matthey for their help in technical parts of this study. The work was supported by Swiss National Science Foundation grant to JvdM 31003A_175638 and by a SystemsX.ch Interdisciplinary grant to CM and JvdM. The funders had no role in study design, data collection and analysis, decision to publish, or preparation of the manuscript.

## Supporting Information Legends

**Figure S1: Protein domain predictions in the ICE*clc* bistability regulators BisR, BisD and BisC**

Regulatory genes BisR (A), BisD (B) and BisC (C) are schematically represented by arrows with indicated size (base-pairs). Corresponding predicted protein domains using Phyre2 (prediction confidence within brackets) are shown below the respective genes, with amino acid positions indicated.

**Figure S2: Gene organization of ICE*clc*-related regulatory loci in different bacterial species**

Diagrams show gene organization (colored arrows, to scale) of the ICE*clc reg* locus (*Pseudomonas knackmussii* B13, Accession number HG322950.1), and homologs in *Pseudomonas aeruginosa* Carb01_63 (CP011317.1), *Herminiimonas arsenicoxydans* (CU207211.1); *Acidovorax* sp. JS42 (CP000539.1); *Burkholderia cenocepacia* ST32 (CP011917.1); *Pseudomonas aeruginosa* HS9 (CP030861.1); *Xanthomonas citri* subsp. *citri* AW13 (CP009031.1); *Pseudomonas aeruginosa* W60856 (CP008864.2); and *Cupriavidus nantongensis* X1 (CP014844.1). Genes of unknown function (dark grey) inserted between *orfs96323* and *95213* in strains AW13, HS9 (2) and W60856 gene regions. Numbers below genes indicate the percentage of amino acid identity and coverage to the one from ICE*clc*. ni: no identity.

**Figure S3: Differential fluorescence expression from a single copy dual reporter in *P. putida*.**

Diagrams show violin distribution plots of individual cell fluorescence of *P. putida* from *P*_*int*_ (eCherry) and *P*_*inR*_ (eGFP), complemented with the indicated plasmids, after 24 h growth in succinate minimal medium and induced with 0.05 mM IPTG. Note different fluorescence intensity scales of eCherry and eGFP, the skewed distributions upon induction with BisDC only, and the differences of plasmids pMEEE6.4-alpA (*reg*), pMEEE6.4 and pMEEE6.4PstI on eCherry and eGFP output.

**Figure S4: Unimodal P_tac_ but bimodal P_int_-expression in *P. putida* with reconstructed ICE*clc* bistability generator.**

(A) Unimodal expression of eCherry from single copy P_tac_-*echerry* chromosomal insertion in *P. putida* upon increasing IPTG concentration. (B) Bimodal eCherry fluorescence from P_int_ in presence of pME-based plasmid carrying the *P_alpA_-inrR* loci, upon induction of BisR from a single copy P_tac_-*bisR* insertion in *P. putida* with increasing IPTG concentrations. Shown are quantile-quantile plots of the observed versus expected distribution of fluorescence levels (n= number of observed cells summed from 10-12 technical replicates).

**Figure S5: Maximum likelihood tree of BisDC homologs**

The evolutionary history was inferred by using the Maximum Likelihood method based on the Tamura-Nei model [65]. The tree with the highest log likelihood (−22463.69) is shown. The percentage of trees in which the associated taxa clustered together is shown next to the branches. BisDC representatives also displayed in Fig. S2 are highlighted. C.nantogensis X1 sequences are artifically linked as they came from two individual genes.

**SI Table 1: Plasmid and used strain numbers of *P. putida* derivatives**

**SI Table 2: Subpopulation size of *P. putida* ICE*clc* cells expressing eCherry/eGFP fluorescence from a Pint-/PinR-double reporter inserted in single copy upon overexpression of key regulators from the transfer competence cascade.**

**SI Table 3: Used primers in the study.**

**SI model**

ICEclc transfer competence bistable regulatory model

REACTIONS

A: BisDC feedback loop on alpA-promoter

B: BisDC feedback loop on alpA-promoter, initiated by BisR

C: TciR activating bisR-promoter

D: BisDC activating three downstream (late) promoters

E: BisDC activating GFP reporter.

**SI code**

ICE stochastic model: code for implementation in Julia

